# Loop modelling provides insights into regulation of mTORC1 activity via DEPTOR dimerisation

**DOI:** 10.1101/2024.03.28.587015

**Authors:** Aik-Hong Teh, Tamao Hisano

## Abstract

mTOR regulates cell growth by forming the mTORC1 and mTORC2 complexes. DEPTOR partially inhibits mTORC1, which in turn phosphorylates and inactivates it. Despite the mTORC1–DEPTOR structures, the exact mechanism remains unclear largely because functionally flexible key elements, DEPTOR’s linker in particular, are unresolved. By taking DEPTOR’s dimerisation into consideration, our modelling of these missing loops suggests that monomeric DEPTOR bound to mTORC1 in a non-inhibitory mode, while the domain-swapped dimeric DEPTOR could interact with mTORC1’s FRB domain and block the kinase’s catalytic site with its linker. These two states indicate that linker phosphorylation inactivates DEPTOR possibly by disrupting its dimerisation, which could tether the linker to the kinase domain to enhance mTORC1 inhibition. In addition to DEPTOR, mTOR’s kα9b–kα10 loop, which harbours the S2481 autophosphorylation site, and mSIN1’s flexible domains in mTORC2 might act as inhibitory elements too.

---

mTOR (mechanistic target of rapamycin) is a master regulator of cell growth conserved from yeast to humans. It is a huge serine/threonine kinase from the PIKK (phosphoinositide 3-kinase-related kinase) family^1^ (Fig. 1) and forms two complexes, mTORC1 and mTORC2, that act in response to different environmental cues. In vertebrates, both complexes additionally interact with DEPTOR (DEP domain-containing mTOR-interacting protein), which inhibits mTORC1^2,3^ but itself is also phosphorylated and inactivated by mTORC1^3^. DEPTOR consists of a long linker sandwiched between two DEP domains, DEP1 and DEP2, and a PDZ domain (Fig. 1). The linker harbours a degron between S286 and S291, SSGYFS, and phosphorylation of the three serine by mTORC1 and the kinase CK1α leads to DEPTOR’s ubiquitylation by SCF^βTrCP^ and subsequent proteasomal degradation^4-6^. As a component of the mTOR signalling pathway, DEPTOR deregulation has been associated with the development of many diseases including cancer^7^.

**Fig. 1.**
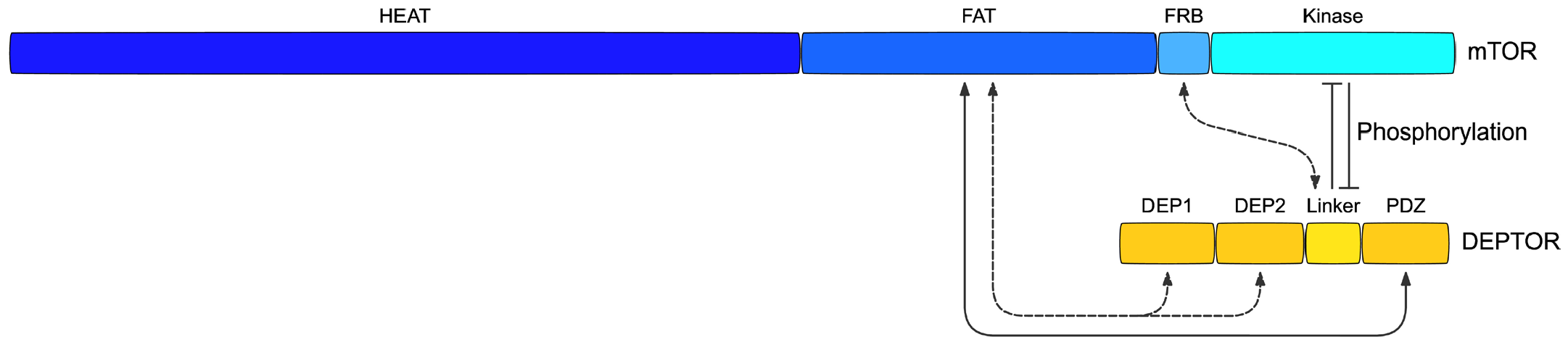
Known interactions between mTOR and DEPTOR. DEPTOR’s PDZ domain binds directly to mTOR’s FAT domain; in addition, either DEPTOR’s DEP1–DEP2 domains also bind to the FAT domain, or its linker interacts with mTOR’s FRB domain. However, it is unclear how unphosphorylated DEPTOR inhibits mTOR’s kinase activity partially, or how phosphorylation of its linker by mTOR inactivates DEPTOR.

The precise mechanism for how DEPTOR regulates mTOR activity is still not fully understood. Two cryo-EM structures of the mTORC1–DEPTOR complex have shed some light on their interaction^3,8^, but not without conflicting results. DEPTOR’s PDZ domain is found to bind to mTOR’s FAT domain in both structures (Fig. 1), while its linker is mostly unresolved. In one of them, DEPTOR binds as a monomer with its DEP1–DEP2 domains additionally interacting with nearby helical repeats of the FAT domain, a binding mode suggested to allosterically suppress mTOR activation^8^. However, the DEP1–DEP2 domains alone reportedly exhibits no inhibition in the other study^3^. Unbound DEPTOR has further been proposed to bind as a substrate at high concentrations to interfere with access to the kinase^8^, but, again, DEPTOR has been shown not to compete as an alternative substrate to inhibit mTORC1 — not only is the DEP1–DEP2–linker variant without the PDZ domain non-inhibitory, but the non-phosphorylatable 13A DEPTOR, with all the 13 serine/threonine in its linker mutated to alanine, also inhibits mTORC1 as well^3^.

DEPTOR’s DEP1–DEP2 domains, meanwhile, are missing altogether in the other cryo-EM structure, while an unidentified linker region is found binding to mTOR’s FRB domain^3^. As mTORC1 inhibition requires minimally the linker–PDZ region, this binding mode offers a straightforward mechanism in which the linker physically blocks access to the kinase^3^. Nevertheless, it is unknown how DEP1–DEP2, whose removal increases DEPTOR’s IC_50_ (half maximal inhibitory concentration) 3.5-fold^3^, acts to enhance mTOR inhibition. The retention of mTORC1 inhibition upon DEP1–DEP2 removal in this study^3^ also contradicts the loss of mTORC1 inhibition upon DEP1’s D120A/D121A mutations, which presumably disrupt the DEP1–FAT binding interface, in the other study^8^. Crucially, both studies do not explain how linker phosphorylation inactivates DEPTOR *in vitro* without proteasomal degradation^3^.

Further adding to the puzzle is the state of DEPTOR’s oligomerisation — DEP1–DEP2 has been characterised and crystallised as both a monomer^8,9^ and a dimer^8,10^. The crystal structure of monomeric DEP1– DEP2 shows a dumbbell conformation^9^, while that of dimeric DEP1–DEP2, notably, harbours a domain-swapped DEP1 which is stabilised by an intermolecular disulphide bond^8^. DEP domains are known to dimerise by domain swapping^11,12^ — the DEP1 domain of the chemotaxis-mediating P-Rex1, which likewise contains two DEP domains and two PDZ domains, forms a highly similar domain-swapped dimer^12^. DEPTOR dimerisation, however, is not taken into account in many studies, which have purified the protein and its variants in the presence of reducing agents such as TCEP^3,8^ or DTT^9^.

Vital details of flexible key elements, in particular DEPTOR’s linker and the unstructured segment between kα9b–kα10 of mTOR’s kinase domain, are also missing in these mTORC1–DEPTOR structures^3,8^. The kα9b– kα10 segment, which plugs one end of mTOR’s catalytic cleft^13^ and harbours the S2481 autophosphorylation site^14-16^, have been found to influence mTORC1 activity too^13,17,18^. These missing elements, however, are difficult to determine structurally via empirical methods due to their inherent flexibility, which is likely important for their functions. Building on these mTOR–DEPTOR complexes, we have attempted an alternative mechanism for the regulation of mTORC1 by DEPTOR by taking the missing elements, through modelling, and DEPTOR dimerisation into consideration. Our models suggest that DEPTOR was active when it bound to mTOR as a dimer and inactive as a monomer, which may represent the two mTORC1–DEPTOR structures respectively^3,8^, and phosphorylation of its linker might inactivate DEPTOR by breaking the dimer apart.

## Results

### Linker of FAT-bound monomeric DEPTOR is unlikely to block mTORC1’s kinase domain

To gain a picture of how mTORC1 and DEPTOR interact, their disordered segments were modelled as extended loops. Modelling of the linker of monomeric DEPTOR, which binds to mTOR’s FAT domain with its DEP1– DEP2 and PDZ domains^8^, was comparatively straightforward (Fig. 2). mTOR’s disordered FAT segment (K1815–Q1866) and its kinase domain’s kα9b–kα10 segment (K2437–E2491), both predicted to be unstructured by AlphaFold (UniProt P42345)^19^, were also constructed. DEPTOR’s modelled linker was barely long enough to reach mTOR’s kinase domain by encompassing mLST8 (Fig. 2), and it was also unable to interact with mTOR’s FRB domain as observed in the FRB-bound complex^3^ nor act across the mTORC1 dimer interface. Crucially, the linker would most likely be prevented from looping over mLST8 by the two mTOR loops as well as mLST8 itself, and wiggle away from mLST8 with higher probability in the absence of any robust interaction between them. DEPTOR therefore did not seem capable of inhibiting mTORC1 in the monomeric form, which might represent an inactive state.

**Fig. 2.**
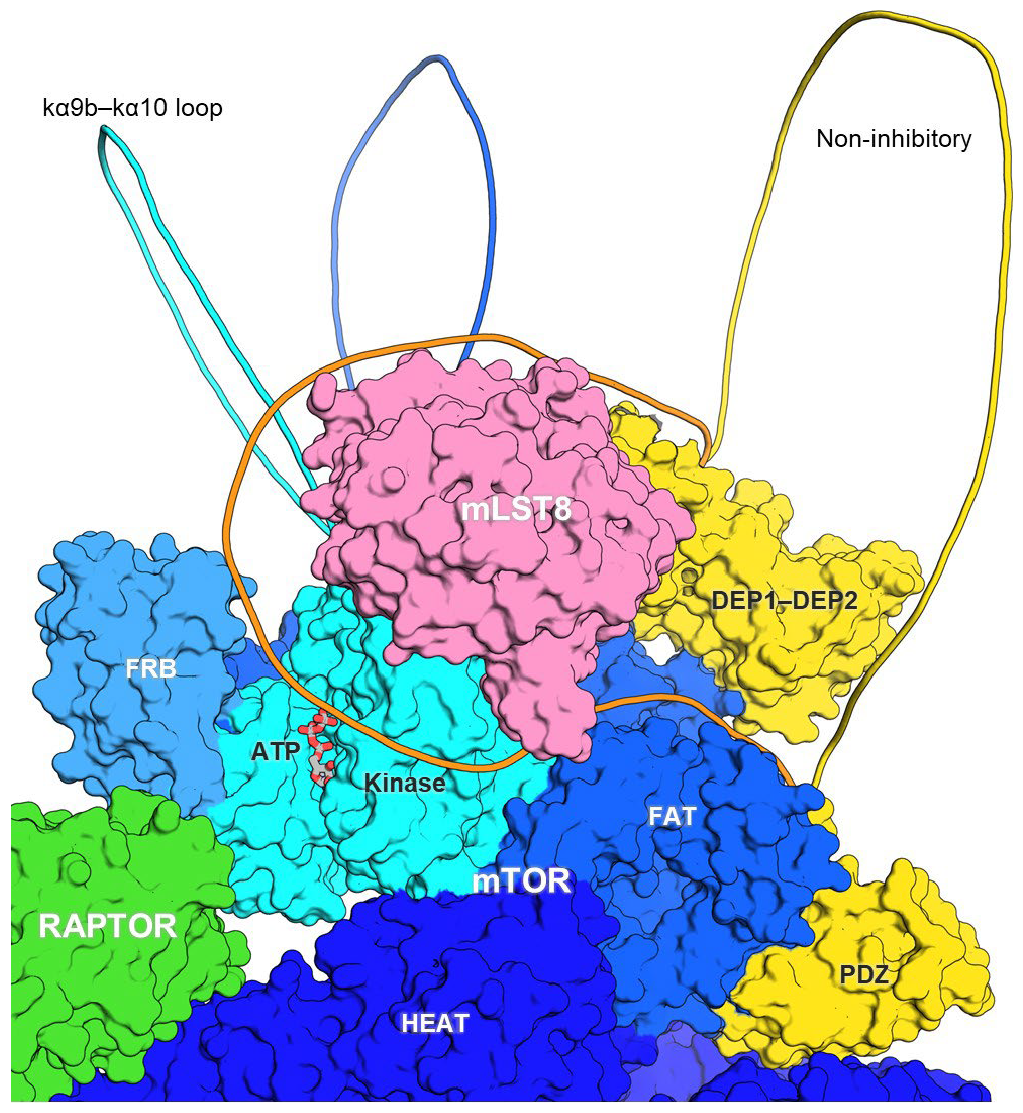
Monomeric DEPTOR is unlikely to bind at mTORC1’s kinase domain. Unstructured regions of the mTORC1–DEPTOR structure were modelled as loops. With both its DEP1–DEP2 and PDZ domains bound to mTOR’s FAT domain, DEPTOR’s linker (orange) could barely encompass mLST8 to get close to mTOR’s kinase domain but was unable to interact with its FRB domain. Since the linker would most likely be prevented by mTOR’s two fluctuating loops, K1815–Q1866 (blue) and the kα9b–kα10 loop (cyan), as well as mLST8 from reaching the kinase domain, the linker would probably wiggle in a non-inhibitory conformation (yellow).

### Linker of dimeric DEPTOR can sterically obstruct mTORC1’s kinase domain

In the FRB-bound complex, DEPTOR’s DEP1–DEP2 is instead missing but a small undefined region of its linker binds to mTOR’s FRB domain^3^. The linker was of sufficient length to be modelled, through S260–V264, to interact with FRB by traversing the N-terminal region of mTOR’s FAT and kinase domains, effectively blocking access to the kinase’s catalytic cleft (Fig. 3a). Surface potential computation showed that, intriguingly, this route was largely electronegative in contrast to the adjacent C-terminal region of the FAT domain which was electropositive (Fig. 3b). This implied that a phosphorylated linker might be repelled from the electronegative region, possibly swinging towards the electropositive region and becoming non-inhibitory (Fig. 3a,b). Since it still inhibits the monomeric mTOR^ΔN^-mLST8 complex^3^, DEPTOR was considered to act on the same mTOR monomer that its PDZ bound to instead of across the mTOR dimer interface. DEPTOR’s missing DEP1–DEP2 could also be modelled as the domain-swapped dimer with an intermolecular C102 disulphide bond^8^ in both inhibitory and non-inhibitory modes (Fig. 3c,d); dimeric DEPTOR was hence able to inhibit mTORC1.

**Fig. 3.**
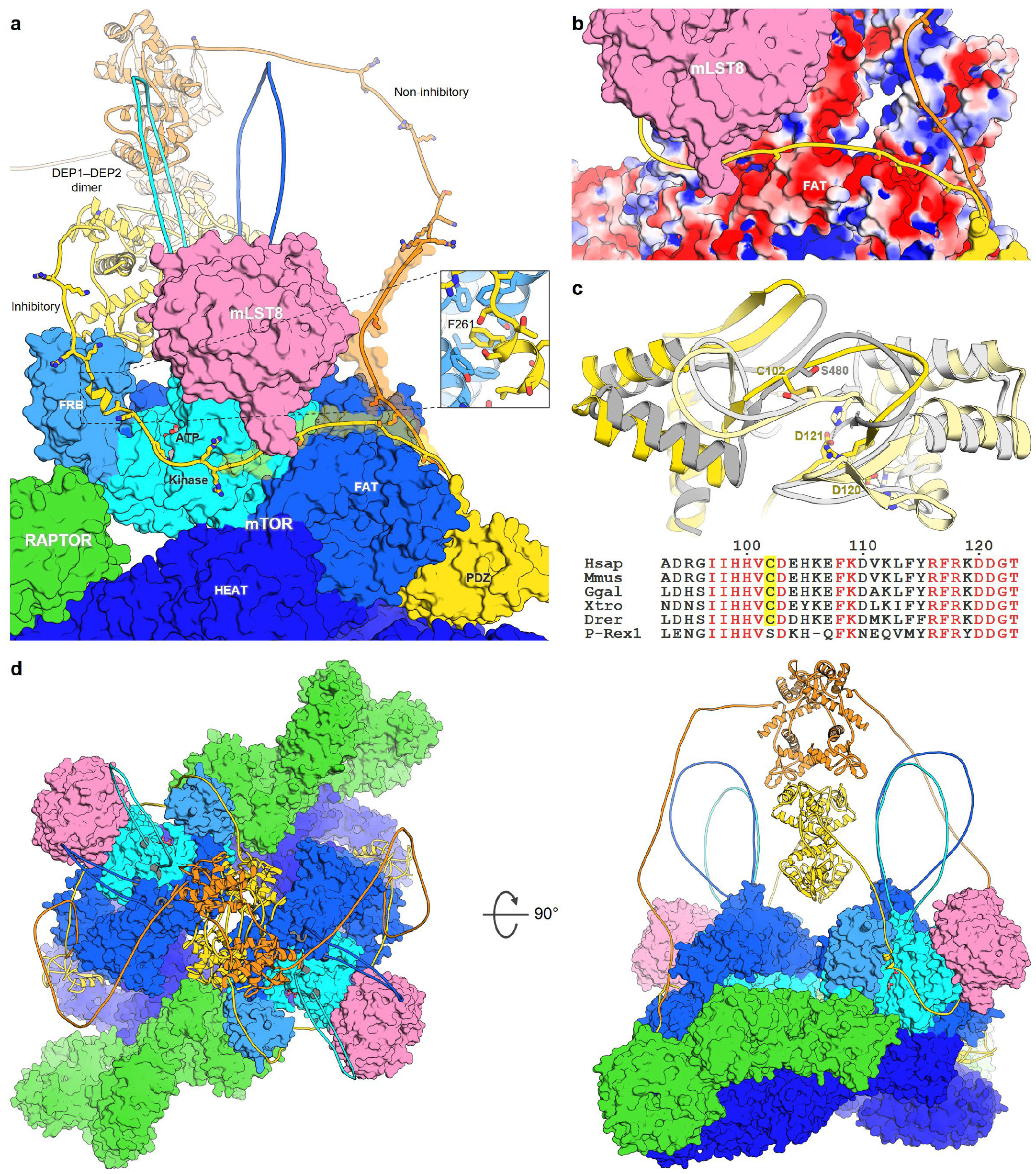
Dimeric DEPTOR partially blocks mTORC1’s kinase domain. (**a**) Without its DEP1–DEP2 domains binding to mTOR’s FAT domain, DEPTOR’s linker was sufficiently long to interact with mTOR’s FRB domain and block its kinase domain (yellow), or swing away into a non-inhibitory conformation (orange). The linker’s basic residues and five mTOR-mediated phosphorylation sites are shown as sticks, while the P box as transparent surfaces. (**b**) DEPTOR’s linker in the inhibitory conformation (yellow) traversed an electronegative FAT region (red) to reach the kinase domain. (**c**) The domain-swapped DEP1 dimers of both DEPTOR (yellow) and P-Rex1 (grey) are structurally highly similar, although DEPTOR’s intermolecular C102 disulphide bond is not conserved in P-Rex1. (**d**) With its DEP1–DEP2 dimer centred onto the mTORC1’s two-fold axis, dimeric DEPTOR was compatible with both the inhibitory (yellow) and non-inhibitory (orange) modes.

### mSIN1 instead of DEPTOR restricts access to mTORC2’s kinase domain

mTORC2’s kinase domain is more crowded compared to mTORC1’s — in addition to RICTOR wrapping around mTOR’s FRB domain, mSIN1 also interacts with both RICTOR and mLST8 around mTORC2’s kinase domain^20^. RICTOR and mSIN1’s disordered segments were similarly built onto the mTORC2–DEPTOR complex^8^ (Fig. 4). Notably, placement of mSIN1’s missing CRIM, RBD and PH domains showed that these three flexible domains could be revolving around the kinase domain, acting to obstruct its catalytic cleft and limit access to only CRIM-recruited substrates. DEPTOR’s linker would thus be prevented by both RICTOR and mSIN1 from interacting with mTOR’s FRB and kinase domains, and would bind only in a non-inhibitory fashion (Fig. 4).

**Fig. 4.**
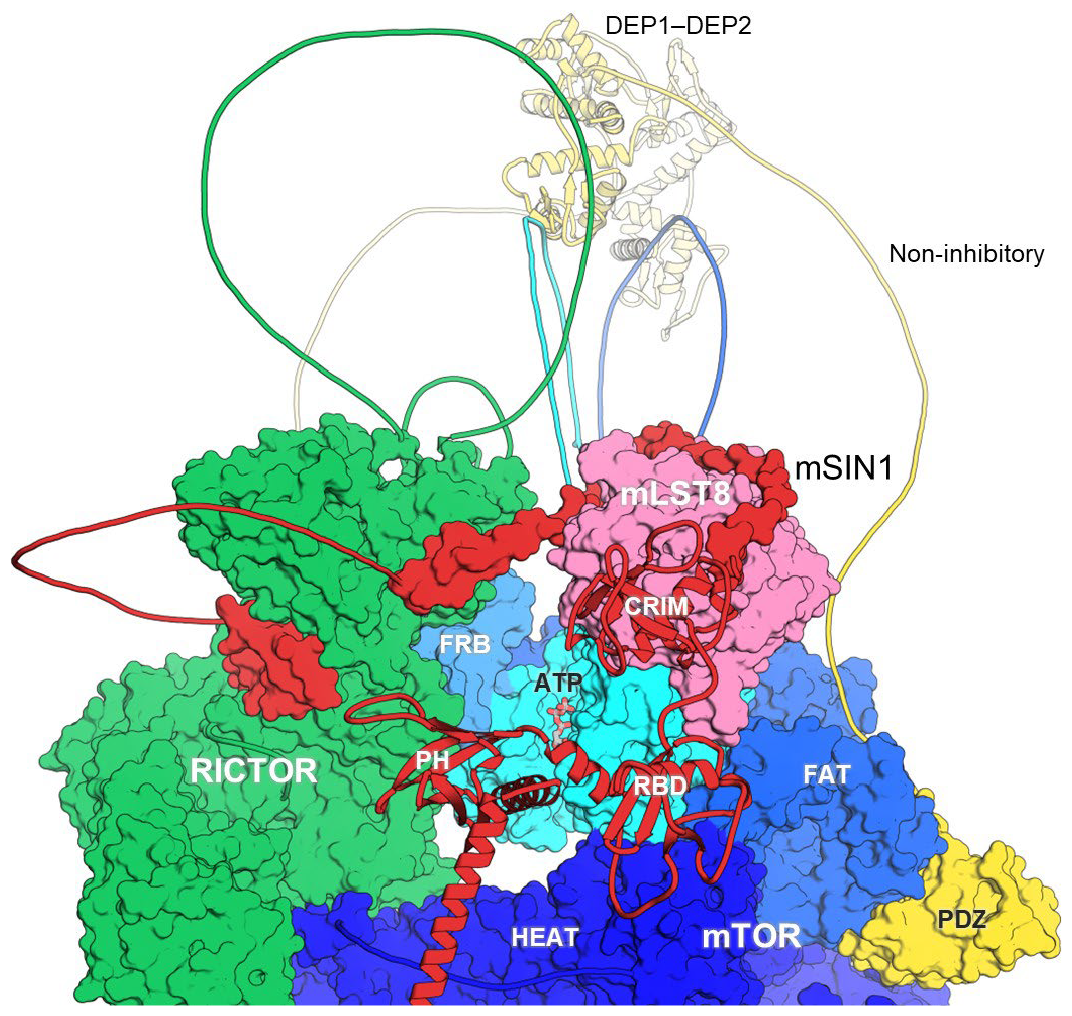
mSIN1 instead of DEPTOR blocks mTORC2’s kinase domain. DEPTOR’s linker likely adopted a non-inhibitory conformation (yellow) as it was prevented from interacting with mTOR’s FRB domain by RICTOR (green) as well as mSIN1 (red), whose flexible CRIM, RBD and PH domains instead could obstruct access to the kinase domain.

### Phosphorylation sites of unbound DEPTOR form a turn to enter mTORC1’s catalytic cleft

Due to length constraints, most of the phosphorylation sites on the linker of mTORC1-bound DEPTOR, particularly those close to the PDZ domain such as S299, were unable to reach mTOR’s kinase domain (Fig. 3a). Moreover, these phosphorylation sites were also in the opposite orientation relative to the peptide substrate bound to SMG1^21^, a structurally related PIKK kinase, and hence could not be acted on by mTOR. Unbound DEPTOR, on the other hand, could be phosphorylated by mTORC1 as its linker could readily access the kinase’s catalytic cleft, which is plugged by the kα9b–kα10 loop at one end^13^, by forming a turn (Fig. 5a). This might explain mTOR’s preference for phosphorylation sites with a succeeding proline at the +1 position^22^, as proline commonly facilitates the formation of turns.

**Fig. 5.**
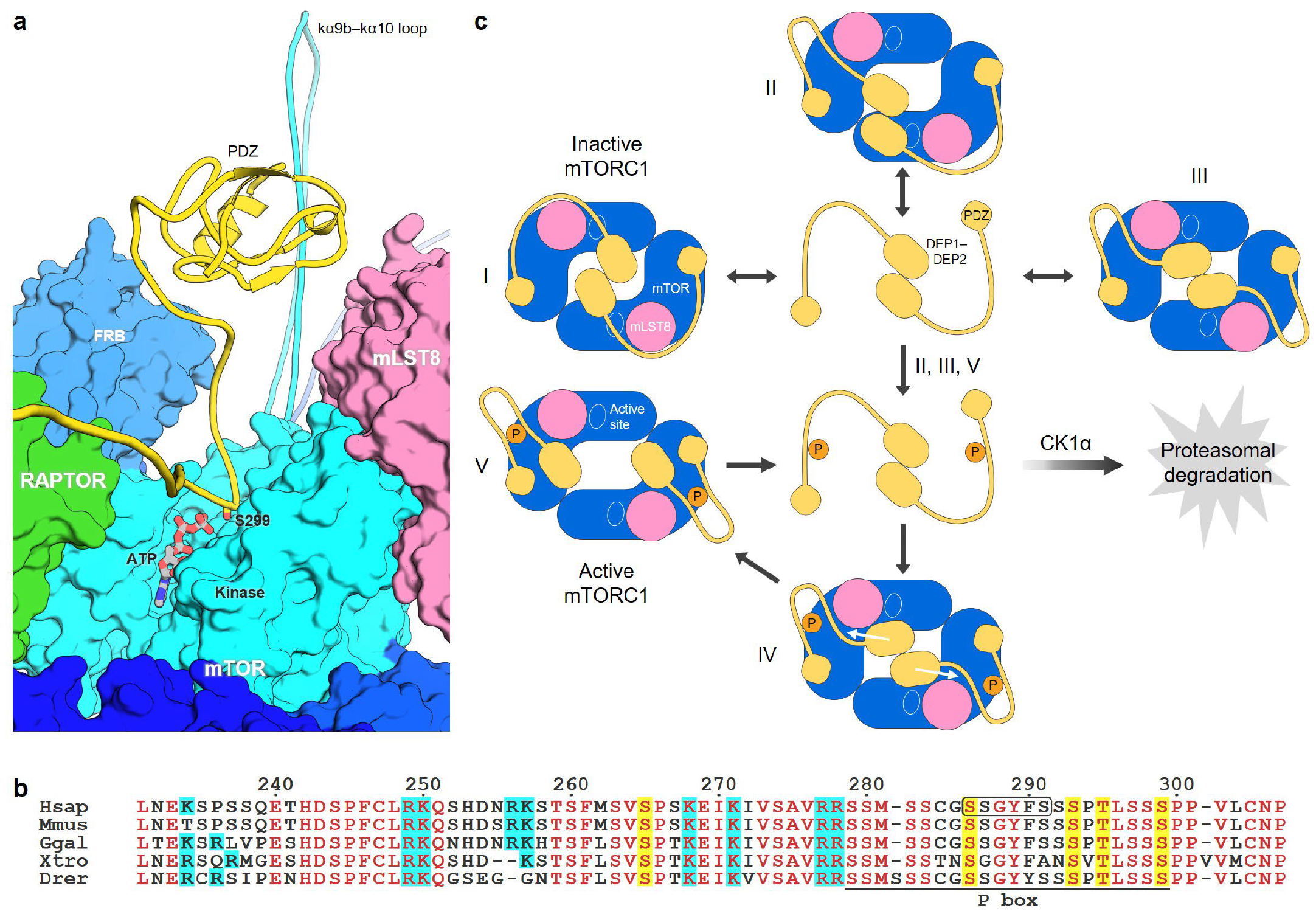
Proposed mechanism for mTORC1–DEPTOR interactions. (**a**) Forming a turn may help the linker of unbound DEPTOR enter mTORC1’s catalytic cleft, which is sealed by the kα9b–kα10 loop at one end. (**b**) Located downstream of several conserved basic residues (cyan), the P box in DEPTOR’s linker contains the degron (boxed), and four of the five serine/threonine (yellow) that are phosphorylated by mTORC1 *in vitro*. (**c**) In the proposed mechanism, unphosphorylated dimeric DEPTOR bound to mTORC1 in three ways (I, II and III), resulting in partial inhibition of mTORC1. Linker phosphorylation fully inactivated DEPTOR by pulling the dimer apart via linker compaction and interactions with mTOR’s FAT domain, subsequently speeding up the phosphorylation of unbound DEPTOR for proteasomal degradation.

## Discussion

Previously we had attempted an mTORC1–DEPTOR model^10^ before the knowledge of their precise interactions was available. At present, despite the two mTORC1–DEPTOR cryo-EM structures^3,8^, how DEPTOR acts to inhibit mTOR still remains largely unclear. These two structures with notable differences — one with DEPTOR’s DEP1–DEP2 domains bound to mTOR’s FAT domain^8^ while the other with DEPTOR’s linker bound to mTOR’s FRB domain^3^ — probably are in fact mutually exclusive, since DEPTOR was unable to bind both ways simultaneously due to length constraints (Fig. 2). With both its DEP1–DEP2 and PDZ domains bound to mTOR’s FAT domain^8^, monomeric DEPTOR was unlikely to inhibit mTORC1 and its linker would likely wiggle away from mTOR’s kinase domain with high probability (Fig. 2). By interacting with mTOR’s FRB domain via an electronegative FAT region, in contrast, DEPTOR’s linker could readily obstruct the kinase domain and inhibit mTOR (Fig. 3a,b). This concurs well with an mTOR inhibition which is dependent on neither substrate identity nor linker phosphorylation, as the 13A DEPTOR is still capable of inhibiting mTORC1^3^. DEPTOR’s DEP1–DEP2 domains, perplexingly, are missing in the FRB-bound structure^3^ — it is unclear why the weak FRB–linker interaction with a *K*_d_ of ∼500 μM^3^ is preferred over the interaction between FAT and DEP1–DEP2 that covers almost 10 nm^2^ of interface^8^.

DEPTOR’s oligomeric state is contentious too — DEP1–DEP2 has notably been characterised as monomeric in the presence of 1 mM TCEP^8^ or DTT^9^, but dimeric in the absence of any reducing agents^10^. Moreover, while the monomeric DEP1–DEP2 purified in DTT also crystallises as a monomer^9^, the one purified in TCEP, intriguingly, crystallises as a dimer containing a swapped DEP1 with an intermolecular disulphide bond^8^. DEP domains are known to form domain-swapped dimers such as the Dishevelled protein DVL2^11^ and the RhoGEF P-Rex1^12^, both of which also switch between the monomeric and dimeric states. In particular, the domain-swapped structure of P-Rex1’s DEP1^12^, which shares 48% identity with DEPTOR’s DEP1, is strikingly similar to the latter’s structure^8^ albeit without the disulphide bond (Fig. 3c). These suggest that DEPTOR may likewise switch between a monomer and a dimer through DEP1 swapping, which may be stabilised by the conserved C102 disulphide bond. Consequently, the presence of 1 mM TCEP during protein purification might have reduced the disulphide bond, while subsequent protein concentration possibly helped DEP1 dimerise again to form the domain-swapped DEP1–DEP2 crystal structure^8^, as well as the FRB-bound cryo-EM structure^3^ in which dimerisation sequestered the missing DEP1–DEP2 from binding to mTOR’s FAT domain. For the mTORC1 structure bound by monomeric DEPTOR^8^, meanwhile, increasing TCEP to 2 mM during sample preparation might have caused dimeric DEP1–DEP2 to dissociate again and bind to FAT. On the other hand, 1 mM DTT, which is superior to TCEP with 100-fold greater thioredoxin reduction due to its smaller size in reaching protein disulphides^23^, could have fully reduced even concentrated DEP1–DEP2 to yield the monomeric crystal structure^9^.

DEP dimerisation has been shown to regulate the activity of DEP-containing proteins — the DEP domain of DVL2, which also contains a PDZ domain and a phosphorylatable linker, dimerises to form an active Wnt signalosome that transduces extracellular signals^11^. To be active, likewise, DEPTOR might need to dimerise so as to prevent its DEP1–DEP2 from binding to mTOR’s FAT domain into a non-inhibitory mode (Fig. 2). DEPTOR has an isoform which lacks DEP1^24^ but may still dimerise since DEP2 (D127–S235) alone is also dimeric^10^; this lends further support to the existence of a dimeric species which may be relevant to DEPTOR’s function. Indeed, deleting DEP1–DEP2 more than triples DEPTOR’s IC_50_ value^3^ — DEP1–DEP2 dimerisation, structurally compatible with the FRB-bound inhibitory mode, could tether the linker to the kinase domain to enhance mTOR inhibition (Fig. 3). It is noted that as the FRB binding is weak^3^, the linker might also swing towards the electropositive FAT region into a non-inhibitory mode (Fig. 3), which may explain why DEPTOR inhibits mTORC1 only partially^3^.

Assuming the two cryo-EM structures^3,8^ represented DEPTOR’s on and off states respectively, a switch may be needed for the on-to-off transition — linker phosphorylation. Under conditions such as insulin deprivation, PRAS40 (proline-rich AKT substrate of 40 kDa), a potent inhibitor with an IC_50_ 16-fold lower than that of DEPTOR^3^, fully inactivates mTORC1 by binding FRB tightly to block the kinase^25-27^. Upon PRAS40 dissociation following mTORC1 activation, DEPTOR’s linker could swing to bind to FRB and start inhibiting mTORC1. However, as the phosphorylation sites of mTORC1-bound DEPTOR were in the reverse orientation relative to the kinase’s catalytic site, only unbound DEPTOR could be phosphorylated. In the presence of substrates such as 4EBP1 (eIF-4E binding protein 1) and S6K1 (S6 kinase 1), which are recruited by RAPTOR via a TOS (TOR signalling) motif^28,29^, DEPTOR would be outcompeted and thus only very little of it was phosphorylated^3^. This is also reflected in the loss of mTORC1 inhibition upon deletion of DEPTOR’s PDZ domain^3^ — only when DEPTOR is anchored to mTORC1 through its PDZ can it compete meaningfully as an inhibitor.

Given that phosphorylated DEPTOR no longer inhibits mTORC1^3^, and that monomeric DEPTOR would probably bind in a non-inhibitory manner (Fig. 2), it is tempting to speculate that linker phosphorylation would convert the active DEPTOR dimer into the inactive monomer. With a *K*_d_ of 7 μM between mTOR’s FAT and DEPOTR’s PDZ^3^, which is comparable to the *K*_M_ of 1.8 μM between RAPTOR and 4EBP1’s TOS motif^27^, unbound phosphorylated DEPTOR would gradually swap places with mTORC1-bound DEPTOR. Phosphorylation is known to affect a protein’s local and global structures by inducing conformational changes, which is especially frequently observed in intrinsically disordered proteins^30^. DEPTOR’s linker, notably, contains several well conserved arginine and lysine residues located upstream a block of conserved phosphorylation sites (S279–S299), hereby dubbed the P box, which harbours the *in vitro* mTORC1 substrates S286, S293, T295 and S299^4^ but totally lacks any basic residue (Fig. 5b). Segregation of oppositely charged residues within an intrinsically disordered sequence is known to induce chain compaction via long-range electrostatic attractions^31^. Therefore, the phosphorylated P box of mTORC1-bound DEPTOR might interact with the upstream basic residues as well as the electropositive region of mTOR’s FAT domain, compacting the linker and anchoring it to FAT (Fig. 5c). Simultaneous linker compaction might eventually pull the DEP1– DEP2 dimer apart, which would quickly be captured by FAT to turn DEPTOR into inactive monomers.

While the proposed mechanism for DEPTOR regulation still remains to be verified, it may help explain certain discrepancies between some studies. Specifically, phosphorylation of the linker’s S293 and S299 has been suggested to promote CK1α-mediated DEPTOR inactivation via proteasomal degradation^4,5^, yet mTORC1 alone is sufficient to inactivate DEPTOR *in vitro*^3^. Mutating the linker’s S286, S293 or S299 to alanine, perplexingly, impedes mTORC1’s ability to phosphorylate and inactivate DEPTOR *in vitro*^4^. One possibility was that phosphorylation of these serine residues might be required for linker compaction; consequently, the mutant DEPTOR dimers could not be pulled apart, so they continued blocking mTORC1 and hence their own phosphorylation. Also, it is baffling that the D120A/D121A mutations in DEP1 result in DEPTOR inactivation^8^, while the DEPTOR mutant without DEP1–DEP2 still inhibits mTOR^3^. As both D120 and D121 are involved in DEP1 swapping^8^ (Fig. 3c), their mutations might have in fact destabilised DEPTOR dimerisation, but the exposed DEP1–DEP2 might still bind mTOR’s FAT domain into the inactive state since their interactions were largely intact.

DEPTOR’s linker, in comparison, seemed unlikely to inhibit mTORC2 because of RICTOR, which wraps around mTOR’s FRB domain^20^, and mSIN1, whose structurally unresolved CRIM, RBD and PH domains would probably revolve around mTOR’s kinase domain and instead act to block it (Fig. 4). The loss of the phosphorylation of the mTORC2 substrate AKT upon truncation of the CRIM domain^32^ but not all the three domains^33^, indeed, substantiates mSIN1’s role as an inhibitor that allows only substrates recruited by the CRIM domain^20,32^, analogous to TOS motifs recruited by RAPTOR^27,34^, to access the kinase domain. Removing CRIM–RBD–PH would thus unblock the kinase domain, possibly allowing mTORC2 to indiscriminately phosphorylate a variety of substrates. Since mTORC2 may phosphorylate DEPTOR too — rapamycin treatment^2^ or RAPTOR knockdown^4^ that disrupts mTORC1 activity still leaves behind a small portion of phosphorylated DEPTOR — either additional interactions might be required for unbound DEPTOR to overcome the mTORC2 inhibition by mSIN1, or the inhibition, like mTORC1 inhibition by DEPTOR, was only partial. These ideas seem to contradict DEPTOR’s role as an mTOR inhibitor, but evidence for the mTORC2– DEPTOR interactions is still lacking and inconclusive, with opposing effects of DEPTOR depletion or overexpression on mTORC1 and mTORC2 activity having been reported^7^.

DEPTOR is believed to inhibit mTORC2 too because its knockdown promotes AKT S473 phosphorylation^2^. However, AKT phosphorylation is found to decrease along with DEPTOR levels in a linear relationship on serum stimulation^2,4,5^. As DEPTOR knockdown may lay open mTORC1’s kinase domain and enable it to also accept AKT — indeed, recombinant mTORC1 is found to cross react with AKT^35^ — an assay without mTORC1 is needed to firmly establish DEPTOR’s action towards mTORC2. On the other hand, DEPTOR overexpression reduces mTORC1 activity but unexpectedly promotes mTORC2 activity, which is attributed to a negative feedback loop arising from mTORC1 inhibition that activates PI3K (phosphatidylinositol 3-kinase) signalling to overcome mTORC2 inhibition by DEPTOR^2^. The PI3K-mediated mTORC2 activation, however, is debatable — PI3K generates PIP_3_ (PI 3,4,5-trisphosphate) that reportedly binds mSIN1’s PH domain to prevent it from interacting with mTOR’s kinase domain^36^, but since removing the PH domain does not improve mTORC2’s activity^33^, it is unsure if the PH domain really binds to and inhibits the kinase domain. Besides mTORC2 possibly being able to pull apart phosphorylated DEPTOR into inactive monomers, it remains to be seen whether DEPTOR also interacts with mTORC2 in any other way, inhibiting or activating.

mTOR’s preference for phosphorylation sites with a succeeding proline probably stems from proline’s readiness to form turns, which facilitate a phosphorylation site’s entry into the kinase’s catalytic cleft that has one end sealed by the kα9b–kα10 loop (K2437–E2491)^13^ (Fig. 5a). As the kα9b–kα10 loop harbours S2481 which is autophosphorylated in both mTORC1 and mTORC2^14^, the loop is able to access the catalytic cleft and thus may act as an inhibitory element too. The antibody mTAb1, directed against mTOR’s D2433–S2450 and found to activate mTORC1^37^, might bind to the kα9b–kα10 loop and prevent it from blocking the catalytic cleft. Similarly, phosphorylated S2481 might restrict the loop’s movement by binding to, for instance, the upstream K2440–R2445 cluster; deleting R2443–G2486 reduces mTOR^ΔN^–mLST8’s kinase activity^13^ possibly because the loop could no longer be restrained by S2481 phosphorylation. It is noted that deleting R2430–S2450^17^ or D2433–A2451^18^ instead activates mTOR, but these deletions also remove part of the kα9b helix (P2425– T2436)^13^ that could have disrupted its conformation and affected the loop’s orientation and movement.

Despite crucial details being missing from the mTORC1–DEPTOR and mTORC2–DEPTOR structures^3,8^, the accumulating knowledge has nevertheless enabled the construction of their possible interactions. Dimeric DEPTOR could physically block mTORC1’s kinase domain with its linker until it was phosphorylated, an event that might break the dimer apart and fully inactivate it. Our proposed mechanism is largely in agreement with mutational studies, while it also implies that DEPTOR may interact with mTORC2 differently. These structural estimations of missing elements highlight the importance of their functional flexibility, and offer fresh insights for further probing the interplay between DEPTOR and the mTOR complexes.

## Methods

### Construction of models

Monomeric DEPTOR’s linker and mTOR’s two unstructured loops (K1815–Q1866 and K2437–E2491) were manually modelled into the mTORC1–DEPTOR cryo-EM structure (PDB: 7PEC)^8^ using *Coot*^38^, and regularised using the geometry minimisation tool of Phenix^39^. For the construction of dimeric DEPTOR, the domain-swapped DEP1–DEP2 dimer (PDB: 7PED)^8^ was docked onto the two-fold axis of the mTORC1 dimer (PDB: 7PEA)^8^ followed by linker modelling, and the FRB–linker interactions were modelled based on the mTOR^ΔN^–mLST8–PRAS40^173–256^ structure (PDB 5WBU)^27^. RICTOR’s and mSIN1’s missing loops were similarly modelled into the mTORC2–DEPTOR structure (PDB: 7PE7)^8^, while mSIN1’s CRIM, RBD and PH domains were docked using its AlphaFold model (UniProt: Q9BPZ7)^19^. DEPTOR’s S299 was modelled onto mTORC1’s catalytic cleft according to the structure of SMG1 in complex with an UPF1 peptide (PDB: 6Z3R)^21^. Surface electrostatic potential was calculated with PBEQ-Solver^40^. All structure figures were generated using PyMOL (https://pymol.org).

## Acknowledgements

This work was supported by the URICAS programme from Universiti Sains Malaysia (1001/PCCB/870036).

## Author contributions

A.H.T. conceived of the project. Both authors performed the modelling and discussed the results. A.H.T. wrote the manuscript with input from T.H.

